# Inter-relationships between changes in stress, mindfulness, and dynamic functional connectivity in response to a social stressor

**DOI:** 10.1101/2020.04.15.040337

**Authors:** James Teng, Stijn A.A. Massar, Julian Lim

**Affiliations:** Centre for Sleep and Cognition, Yong Loo Lin School of Medicine, National University of Singapore, Singapore

**Keywords:** Stress, mindfulness, arousal, cortisol, dynamic functional connectivity

## Abstract

We conducted a study to understand how dynamic functional brain connectivity contributes to the moderating effect of trait mindfulness on the stress response. 40 participants provided subjective reports of stress, cortisol assays, and functional MRI before and after undergoing a social stressor. Self-reported trait mindfulness was also collected. Experiencing stress led to significant decreases in the prevalence of a connectivity state previously associated with mindfulness, but no changes in two connectivity states with prior links to arousal. Connectivity did not return to baseline 30 minutes after stress. Higher trait mindfulness was associated with attenuated affective and neuroendocrine stress response, and smaller decreases in the mindfulness-related connectivity state. In contrast, we found no association between affective response and functional connectivity. Taken together, these data allow us to construct a preliminary brain-behaviour model of how mindfulness dampens stress reactivity.

## Introduction

Facing a stressor triggers a complex cascade of physiological and psychological reactions that prepares the body to respond to physical threat. In social settings however, the effects of these reactions are largely undesirable, for example, having to deal with anxious thoughts and a racing heart while giving an important presentation. Using mindfulness – the practice of focusing one’s attention on the present moment while maintaining a non-judgmental and non-reactive stance to the experiences in it – is an effective strategy for coping with, and dampening the effects of stress in such situations. While the psychological and biological changes associated with mindfulness and stress have been well studied separately, a unified model of mind, brain, and endocrine responses to their interaction has yet to be constructed. Here, we make a first attempt to connect some of these disparate pieces using cortisol and functional MRI connectivity changes resulting from the Trier Social Stress Test (TSST) (Kirschbaum et al., 1993), a widely used laboratory procedure to induce stress (Figure 1).

**Figure 1.**
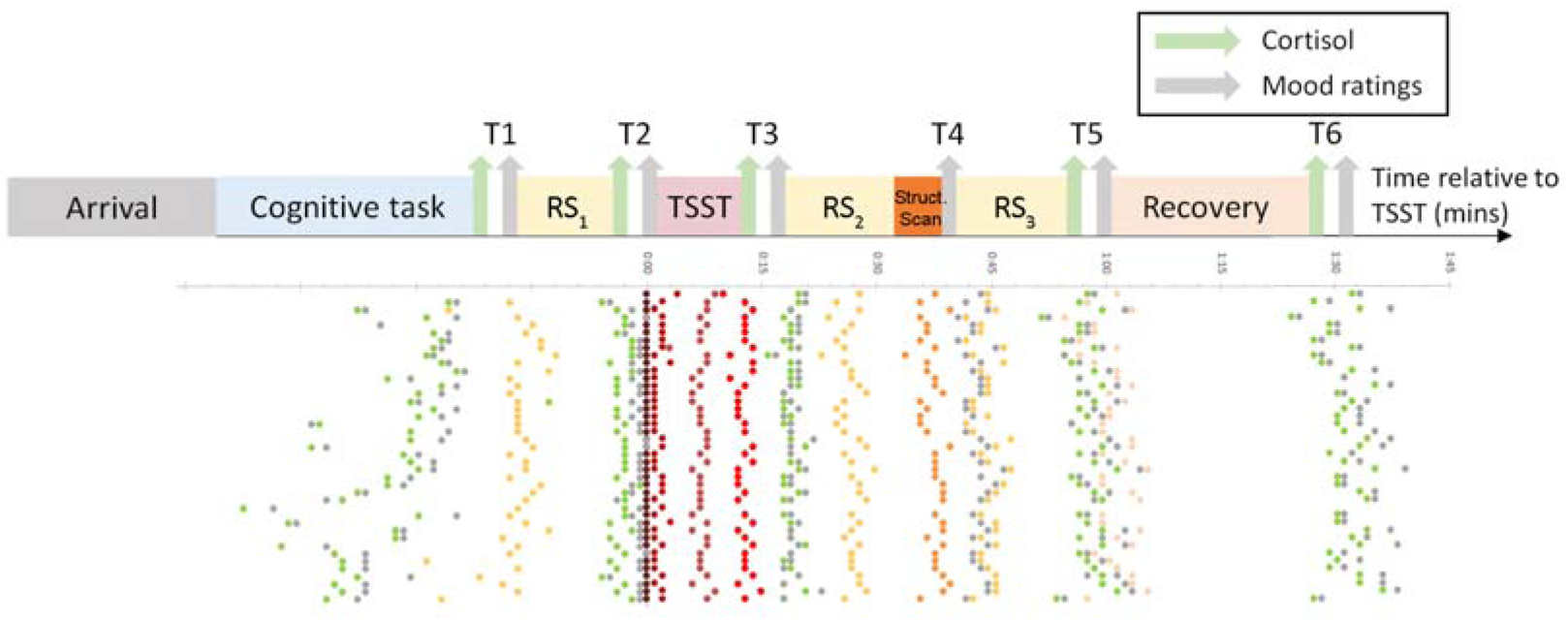
Schematic diagram of experimental protocol. Cortisol and subjective mood ratings were collected at 6 time points, and the main section of the protocol consisted of 3 resting-state fMRI scans, with administration of the Trier Social Stress Test (TSST) between scans 1 and 2. Each row of dots represents a single participant, and the time that they began each stage of the procedure relative to T1.

Undergoing the TSST causes transient but large increases in perceived stress, arousal, and cortisol release (Dickerson & Kemeny, 2004; Kudielka et al., 2007). In addition, there is a growing body of evidence that acute stress also induces changes in resting-state functional connectivity. Exposure to social stress has been linked to increased connectivity between the default mode network and the amygdala (Veer et al., 2011), and more generally to the salience network, while connectivity within the default mode network itself is reduced (Zhang et al., 2019). Recent work has revealed that undergoing the TSST increases activity in hippocampal subregions, which predicts subsequent increases in brain areas related to emotion processing such as the insula (Chang and Yu, 2019). Finally, using graph analysis, Reinelt et al. (2019) showed that stress increases the centrality of the thalamus, and that this increase persists for upwards of 105 minutes after the stressful event

Thus far, little attention has been given to studying the effects of stress on time-varying or dynamic functional connectivity. While traditional static functional connectivity measures provide an estimate of network configuration averaged over a full duration of a scan run, dynamic functional connectivity can identify moment-to-moment changes in network configuration in the order of several seconds (Allen et al., 2014). Detecting such faster changes in connectivity states may be particularly relevant in transient conditions such as acute stress.

In prior work, we used dynamic functional connectivity analysis to identify two connectivity states related to high and low arousal (Wang et al., 2016; Patanaik et al., 2018; Teng et al., 2019). A high arousal state was characterized by high intra-network connectivity, particularly in control areas, and strong anti-correlation between multiple task-positive networks and the default mode network, while a low arousal state was characterized by lower intra and inter-network connectivity. These states fluctuated on a seconds to minutes timescale, and were related to eye closure and attention performance after sleep deprivation (Wang et al. 2016). Later studies showed that these states were systematically and consistently modulated by this manipulation. Based on these findings, we reasoned that the increased arousal caused by stress would result in the reverse effect on functional connectivity than sleep deprivation. We therefore hypothesised that acute stress would cause an increase in time spent in the high arousal state, and a decrease in time spent in the low arousal state.

Mindfulness is the second psychological focus of the current study. This concept was introduced as a tool in psychotherapy in the 1980s (Kabat-Zinn, 1991), and is now an empirically supported method to mitigate the effects of chronic stress (Khoury et al., 2015). It has subsequently been discovered that even in the absence of training, individuals differ in their levels of dispositional mindfulness (Brown and Ryan, 2003), and this natural variation also predicts how one will respond to stress. For example, Brown et al. (2012) demonstrated that high self-reported mindfulness was associated with dampened affective and cortisol response to the TSST, a finding recently reproduced by our own group (Lin et al., in revision).

Researchers have also used resting-state functional connectivity to study the correlates of naturally varying trait mindfulness (Bilevicius et al., 2018; Parkinson et al., 2019) and the effects of mindfulness-based training (Kilpatrick et al., 2011; Doll et al., 2015; Gotink et al., 2016). While results from these experiments are mixed, there is some agreement that mindfulness is associated with increased connectivity within the default mode network, and in particular between the posterior cingulate cortex and ventromedial prefrontal cortex, greater anti-correlations between the default mode network and attentional/control areas, and an increase in connectivity between regions of the ventral attentional network (particularly the insula) and executive control areas (see Mooneyham et al., 2016 for a review). Furthermore, studies have also found variation in dynamic connectivity that corresponds with trait mindfulness (Mooneyham et al., 2017; Lim et al., 2018; Marusak et al., 2018). In a recent study, we identified a mindfulness-related connectivity state (which we named the “task-ready” state) that recapitulates some of the most robust features of findings from static connectivity; high levels of intra-network connectivity in the default mode network and ventral attention network, and strong anti-correlations between these same networks (Lim et al., 2018).

In spite of these intriguing inter-relationships, a unified study of mindfulness, stress, and functional connectivity has not yet emerged. To fill this knowledge gap, we designed an experiment to address the broad questions of i) how dynamic functional connectivity in resting-state fMRI changes following a stressor, and ii) how trait mindfulness moderates this response. We tested several specific hypotheses to answer these questions. Our first prediction, primarily communicated in a separate report (Lin et al., in revision), was that high trait mindfulness would be associated with smaller increases in self-reported stress and cortisol concentration in response to a stressor. Our data support this hypothesis. Our second prediction, as stated above, is that the time spent in arousal-related states will be changed accordingly by stress. Third, we predicted that trait mindfulness would correlate with the proportion of time spent in a third connectivity state, the task-ready state (Lim et al., 2018), and that higher levels of this state at baseline would in turn be associated with a smaller stress response.

## Results

### Subjective stress and cortisol increase following a social stressor

Results from the behavioural ratings and cortisol assays have been reported in a separate communication (Lin et al., in revision). Relevant to this report, we conducted post-hoc t-tests on self-reported stress and salivary cortisol measured before and after the TSST (following ANOVA conducted on data from all 6 data acquisition points). As expected, we observed significant increases in both of these measures (both p < .001 in post-hoc comparisons) after TSST administration, clearly indicating that participants experienced elevated stress following the TSST. In contrast, alertness between pre-TSST and post-TSST did not show a significant increase (5.80 (1.76) to 6.33 (1.69), p = .24 in post-hoc comparison). Full time courses of self-reported stress and salivary cortisol levels across the paradigm are plotted in Figure 2.

**Figure 2.**
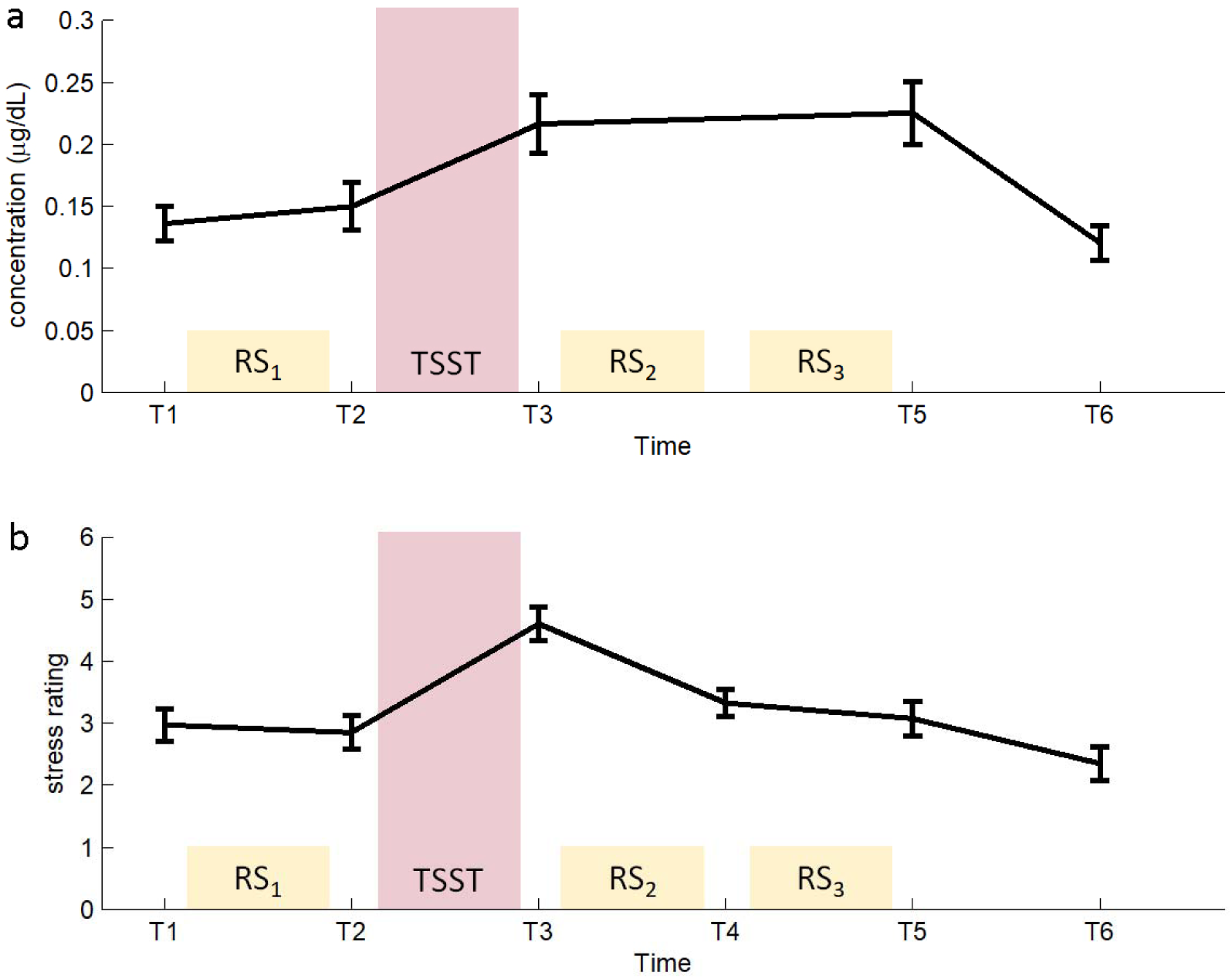
Time courses of (a) cortisol concentration and (b) self-reported stress over the experimental protocol. Both variables increase significantly (p < .001) as a result of the TSST, with perceived stress showing a faster time course of recovery.

### Trait mindfulness attenuates the stress response

Our second key communication in Lin et al. (in revision) was that trait mindfulness was correlated with both the change in self-reported stress (r = −0.41, p = .009) and total salivary cortisol (AUCg; r = −0.38, p = 0.02) across the TSST. In other words, individuals with higher levels of mindfulness experienced an attenuated stress response when confronted with a social stressor.

### Reproducibility of dynamic connectivity states

In a previous dynamic functional connectivity analysis using a similar strategy (Teng et al., 2019), we reported that specifying a five-cluster solution during the classification step yields a set of reproducible connectivity states. To revalidate this finding, we visually inspected the five states in this dataset for their best matching counterpart reported previously, and computed Spearman correlations between the pairs of states. As before, we found high correlations between the centroid pairs (all rho > .82). Importantly, we were able to reproduce the two states associated with high and low arousal in several previous reports (see Figure 3a middle panel: high arousal state; right panel: low arousal state; Patanaik et al., 2018; Teng et al., 2019). The proportion of time spent in these arousal-related states was extracted and used as primary measures in subsequent analysis. We also calculated this variable for a third named state, the “task-ready state” (TRS; see Figure 3a left panel). In a previous study (Lim et al., 2018), we showed that high trait mindfulness was associated with a greater proportion of time being spent in the TRS.

**Figure 3.**
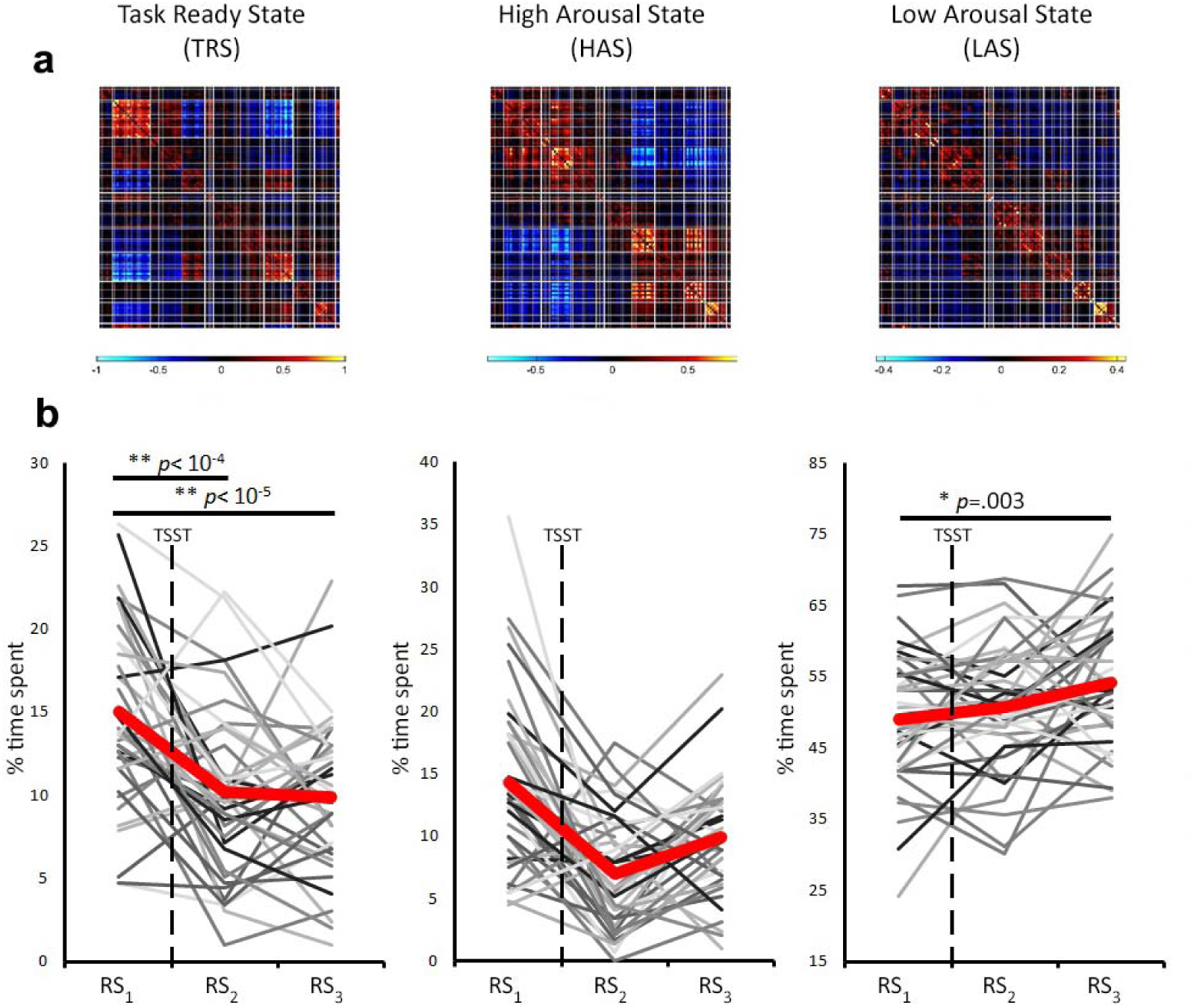
(A) Connectivity centroids of the task-ready state (TRS), high arousal state (HAS) and low arousal state (LAS). (B) Individual trajectories and grand means of changes in the states over the three resting-state scans. Significant differences were found between RS_1_ and RS_2_ for the TRS, and between RS_1_ and RS_3_ for the TRS and LAS.

In the centroid correlation matrix (Supplementary Figure 1a), we observed that there was not a clear one-to-one statistical correspondence between the states obtained in Lim et al. 2018, and those found in this experiment. As a further analysis to determine the distinctness of the connectivity states in our current dataset, we performed dynamic functional connectivity analysis using the pipeline described in this paper on a larger dataset of N = 173 resting-state of fMRI scans collected across our research center. We reasoned that the centroids obtained from this larger dataset would be more stable estimates of the five states. We performed Spearman’s correlations between the canonical centroids derived from that analysis with those from the current dataset, and here showed the desired one-to-one correspondence (Supplementary Figure 1b; all rho > .92), supporting the idea that these states are robust and reproducible. The canonical centroids derived from the larger dataset are freely available as a resource on (https://github.com/awakelab/Dynamic-Functional-Connectivity-MTD-).

### Proportion of time spent in the task-ready state decreases following acute stress

We performed paired-samples t-tests on the three named states between RS_1_ (pre-stress) and RS_2_ and between RS_2_ and RS_3_ (both post-stress) to test whether any of these changed as a result of stress or recovery from stress respectively. Between RS_1_ and RS_2_ (i.e. as a result of performing the TSST), we observed numerical but non-significant increases in both the HAS and LAS (Figure 3b middle & right panels), and a significant reduction (t = 4.76, p < .001) in the TRS (Figure 3b left panel). There were no significant changes in proportion of time spent in any of the states between RS_2_ and RS_3_. Between the two distal scans (RS_1_ and RS_3_), the reduction in TRS was also significant (t = 4.97, p < .001), and a significant increase was seen in LAS (t = 3.21, p = .003).

### Dynamic connectivity changes correlate with changes in cortisol concentration, but not self-reported stress

To test our second hypothesis, we conducted bivariate correlations between the change in self-reported stress and alertness, and ΔHAS and ΔLAS between RS_1_ and RS_2_. None of these correlations were significant (all p > .05).

We next tested for associations between changes in objective stress (cortisol) by correlating total cortisol (AUCg) and cortisol increase (AUCi) over the TSST with ΔHAS, ΔLAS, and ΔTRS over that same period. We found a significant positive correlation between ΔHAS and cortisol AUCi (r = .39, p = .01) and a significant negative correlation between ΔTRS and cortisol AUCg (r = -.33, p = .04) (Figure 4).

**Figure 4.**
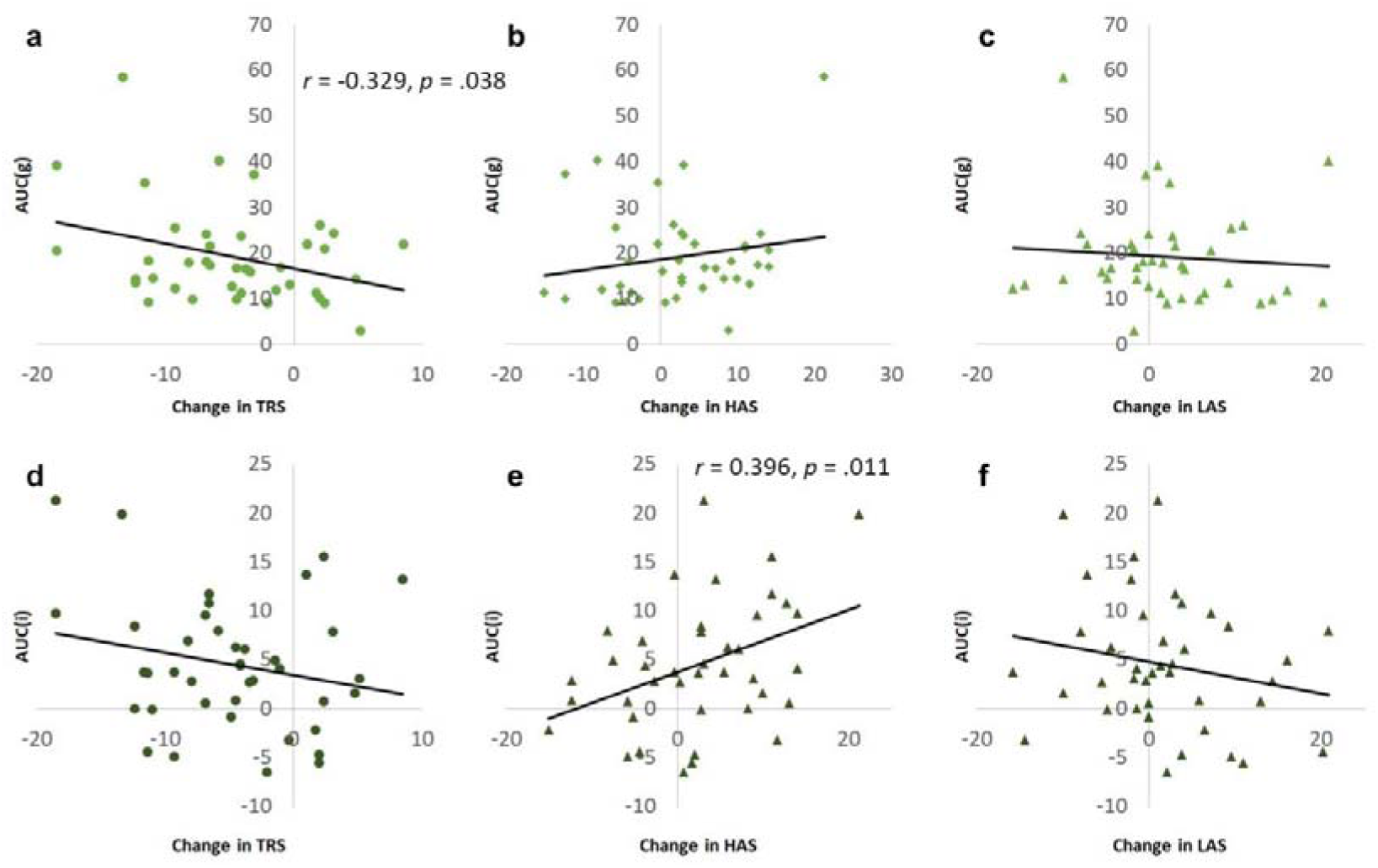
Correlations between cortisol concentration and the change in proportion of time spent in the task-ready state (TRS), high arousal state (HAS) and low arousal state (LAS). Top row: area under the curve of cortisol with respect to ground (AUCg); bottom row: area under the curve of cortisol increase due to stress (AUCi). Significant correlations were observed between ΔHAS and AUCi, and between ΔTRS and AUCg.

### High and Low Arousal States show coupling in two datasets

We tested for coupling among the three named states by correlating ΔHAS, ΔLAS, and ΔTRS across the TSST. Only one of these relationships was significant: we observed a negative relationship between ΔHAS and ΔLAS over this period (Figure 5a; r = -.50, p = < 001). For comparison, we performed a similar analysis on data from our previous report (Teng et al,. 2019), a within-subjects design with fMRI scans obtained in subjects who had undergone 24 hours of total sleep deprivation (compared with when they were well-rested). Again, ΔHAS and ΔLAS between the sleep-deprived and well-rested scans were tightly coupled (Figure 5b; r = −0.64, p < .001).

**Figure 5.**
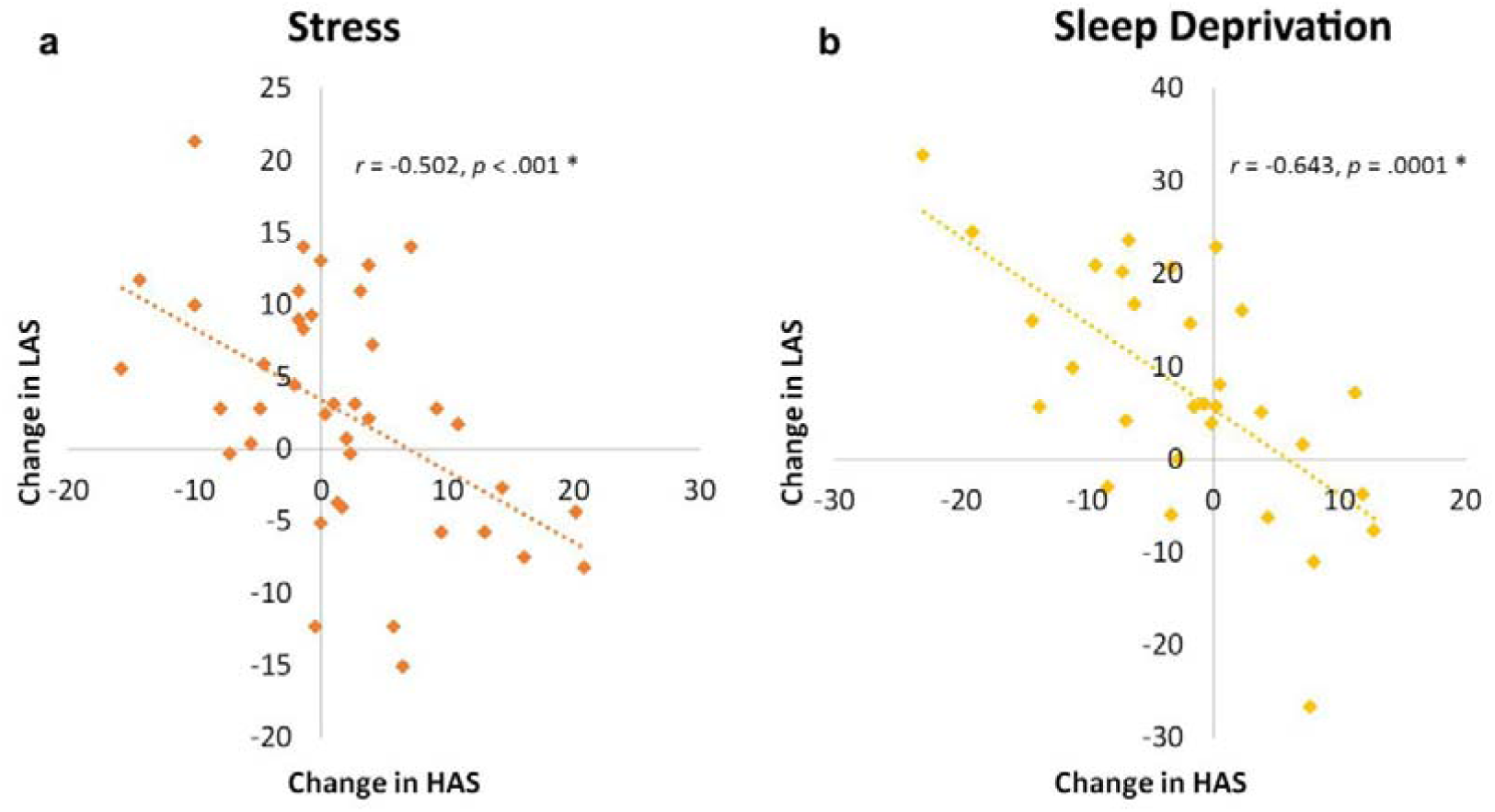
The high arousal state (HAS) and low arousal state (LAS) are coupled and change in tandem across state in two different datasets, a) due to stress, and b) as a result of 24 h of total sleep deprivation.

### Decrease in Task Ready State correlates with trait mindfulness

Given our previous findings linking the TRS with trait mindfulness (Lim et al., 2018), we conducted further analysis to test if a similar relationship held in this dataset. Contrary to our *a priori* hypothesis, we did not find a correlation between trait mindfulness and TRS at baseline (i.e. in RS_1_) (r = -.24, p = .13). Instead, we found that trait mindfulness was significantly correlated with the change in TRS from RS_1_ to RS_2_ (r = .32, p = .045), with greater trait mindfulness associated with a smaller decrease in time spent in this state. This relationship did not hold true for either of the arousal-related states (Figure 6).

**Figure 6.**
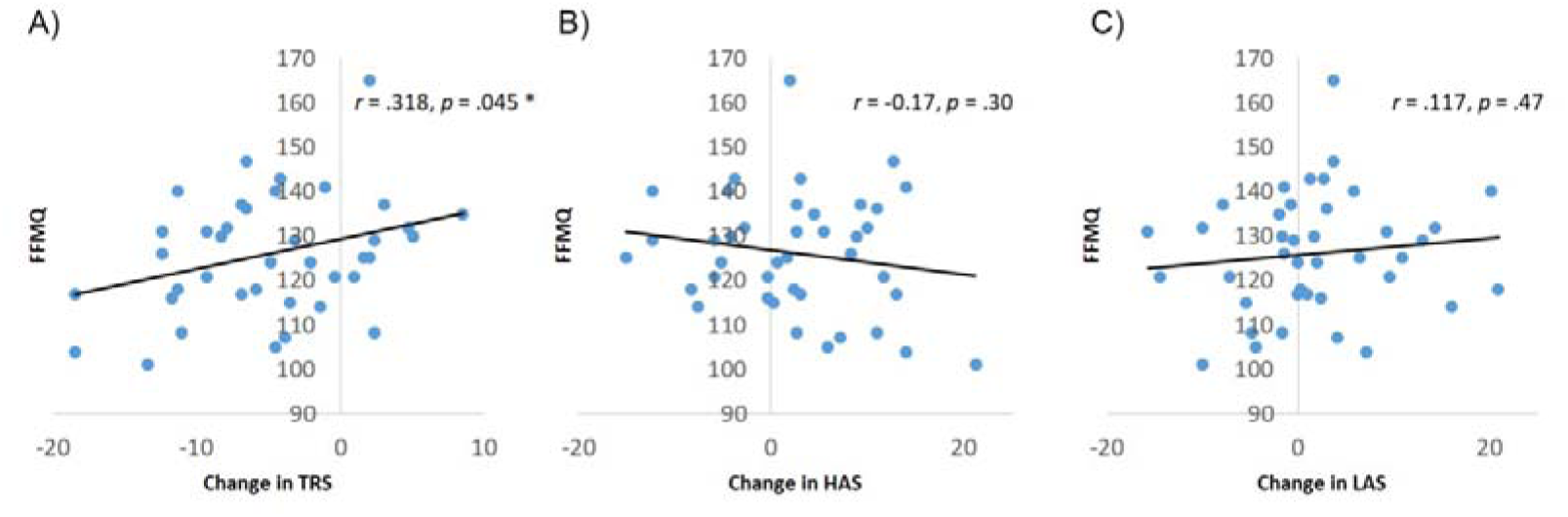
Trait mindfulness correlates positively with the change in proportion of time spent in the task ready state (TRS) due to stress (A), but is not associated with the arousal-related states (B-C).

To illustrate the interrelationship between self-reported data, cortisol, and dynamic connectivity states, we have summarised our findings in Figure 7.

**Figure 7.**
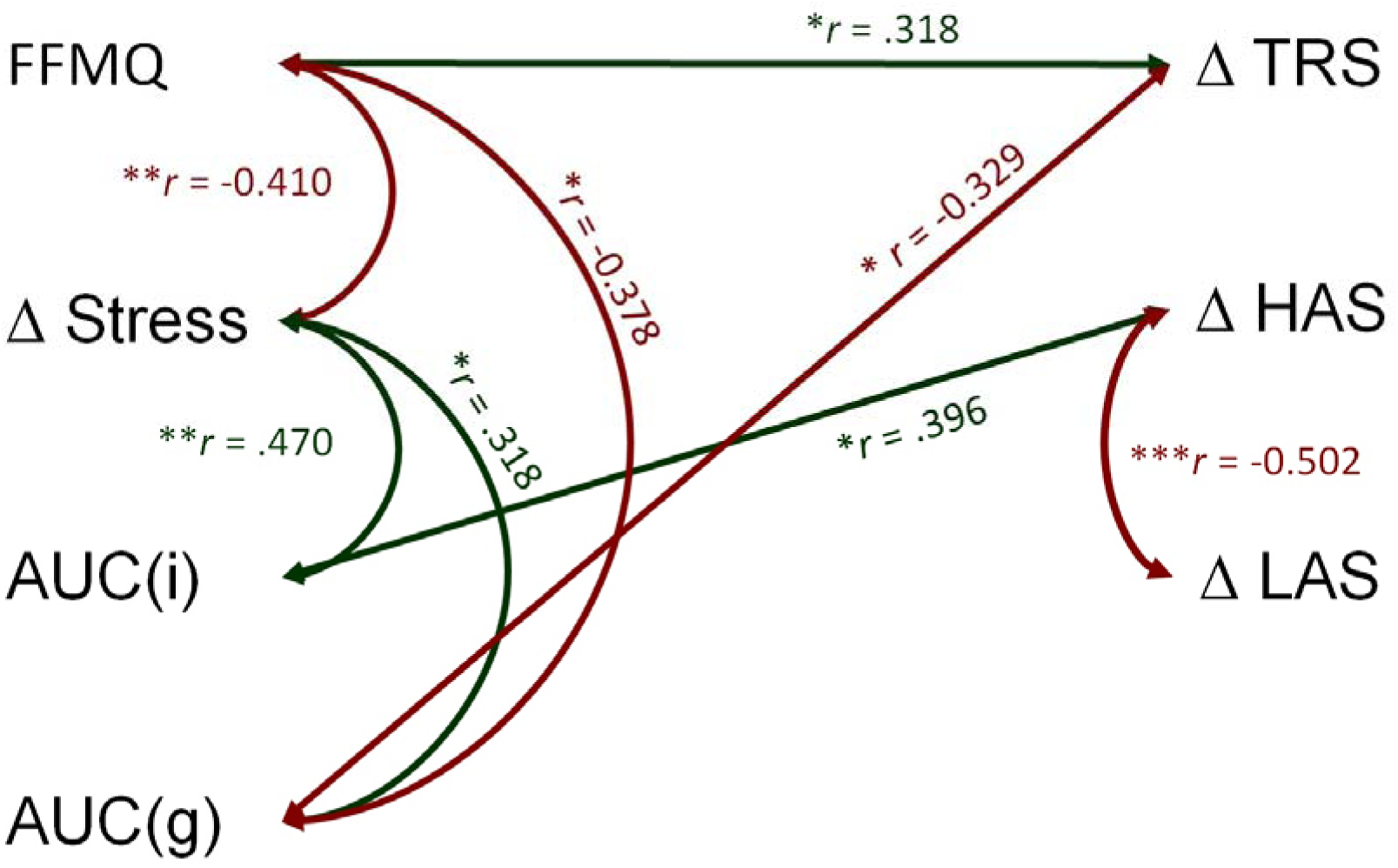
Significant relationships and correlation coefficients among variables of interest. FFMQ = Five facet mindfulness questionnaire, AUCi = increase in area under the curve (cortisol), AUCg = area under the curve with respect to ground (cortisol), TRS = task-ready state, HAS = high arousal state, LAS = low arousal state. * *p* < .05, ** *p* < .01, *** *p* < .001

## Discussion

The current study aimed to link several disparate findings on stress, mindfulness, and their respective effects on functional connectivity. In particular, we were motivated by previous experiments from our group demonstrating the robust association between changes in tonic levels of vigilance and two specific dynamic connectivity states, the high and low arousal states (Wang et al., 2016; Patanaik et al., 2018; Teng et al., 2019), and showing that another dynamic connectivity state, the “task-ready” state, is more prevalent in individuals who are high on trait mindfulness (Lim et al., 2018). As a follow-up to these studies, we reasoned that the preponderance of the HAS and LAS should also change with the heightened arousal caused by increased stress, and should be modulated by trait mindfulness respectively. Contrary to this hypothesis, we did not observe any immediate change in time spent in HAS/LAS, despite the fact that participants clearly demonstrated increased levels of cortisol and subjectively rated stress. Instead, we found large and significant reductions in the TRS as a result of the TSST that were correlated with self-reported trait mindfulness. We then demonstrated that ΔTRS and ΔHAS across the TSST were associated with cortisol reactivity, and followed a similar trajectory to this objective marker, with no recovery to baseline 30 minutes after the stressor.

### Effects of stress on arousal-related states

Thus far, the strongest behavioural link to time-varying connectivity has been with fluctuations in consciousness and arousal. This was first observed in macaques (Barttfeld et al., 2014), which show a decreasing range of functional configurations under increasing levels of sedation by anaesthesia. Later, and also in macaques, Chang et al. (2016) demonstrated that a template-based method could be used to track electrophysiological and behavioural arousal (measured using eyelid closures), and that this method can also track vigilance fluctuations during resting-state fMRI scans in humans (Falahpour et al., 2018).

Using an approach aimed at identifying discrete dynamic connectivity states, Wang et al., (2016) exploited the increased variability of eyelid closures under conditions of sleep deprivation, and showed that periods of higher vigilance were associated with proportionally more time spent in an integrated, “high-arousal” state. This study prompted us to conduct a series of follow-up experiments to test the specificity and reproducibility of the HAS/LAS/behavioural arousal link. Across two further studies, we showed that the HAS and LAS are independently reproducible, that they predict vulnerability to being sleep deprived while participants are still in a well-rested state (Patanaik et al., 2018), and that changes in the proportions of these states following a single night of total sleep deprivation are correlated with a behavioural measure of vigilance (Teng et al., 2019). Finally, the vigilance declines from sleep deprivation were not associated with changes in TRS in this last study, suggesting that there is a dissociation of function between the arousal-related states and TRS.

To date, evidence for the functional significance of the arousal-related states has been generated using sleep deprivation, or reducing levels of arousal, to induce changes in dynamic connectivity. In the current study, one of our primary aims was to test whether the reverse is true: does increasing arousal (by inducing stress) induce the opposite effect to our previous studies, that is, increased HAS and decreased LAS? Surprisingly, we did not find evidence in support of this hypothesis; both the HAS and LAS showed a numerical but non-significant increase following the TSST (with the LAS increase reaching significance in RS_3_), and changes in HAS and LAS were not correlated with changes in self-reported stress or alertness. Increases in HAS were however correlated with cortisol AUCi. Overall, this pattern of data suggests that HAS and LAS responded in some meaningful way to the increase in stress, but that our previous interpretation that they merely reflect arousal levels was overly simplistic.

One alternative explanation for the data are that the high and low arousal state are related more directly to sustained attention than to arousal *per se*. in prior studies, we defined and operationalized arousal using reaction speed on the Psychomotor Vigilance Test (Dinges, 1995), a commonly used measure of sustained attention in human factors and sleep research. While the ability to sustain attention is partially mediated by arousal, the two constructs are not synonymous; arousal is regulated via ascending noradrenergic projections from the brainstem to the thalamus, while sustained attention relies at least in part on top-down, thalamo-cortical circuits (Posner and Petersen, 1990; Sarter et al., 2001). Given our pattern of findings, it is possible that the HAS and LAS index top-down readiness to respond, but not tonic, bottom-up arousal. In support of this possibility, we note that many of the strongly connected edges in the high arousal state centroid originate from the executive control network as well as areas in the dorsal attentional stream. As top-down attention can be adversely affected by both underarousal and overarousal (in an inverted-U function), this could explain the increase in LAS as a result of both stress and sleep deprivation. However, as we did not directly measure sustained attention in this experiment, this interpretation remains speculative.

Another possible interpretation of the results is that increasing arousal via stress may not lie on the same dimension as dampening arousal using sleep deprivation. While noradrenergic activation changes in opposite directions during stress (increase) and sleep deprivation (decrease), cortisol is found to increase both after stress and after sleep deprivation (Spiegel et al., 1999). In fact, sleep deprivation is often considered as a form of physiological stress (McEwen, 2006). In this light it is relevant to note that insufficient sleep can lead to increased cortisol reactivity to the Trier Social Stress Test (Minkel et al., 2014; Massar et al., 2017). In the current study, higher cortisol output after stress was associated with reduction in TRS, but not with any changes in arousal-related states (HAS/LAS). It should be noted that in our previous study sleep deprivation similarly lead to a decrease in TRS (Teng et al. 2019). It is therefore possible that decreases in TRS after sleep deprivation are also related to increased cortisol, while increase in LAS may be more related to reduced noradrenergic tone. This possibility however would still need to be verified.

### The role of trait mindfulness

The primary behavioural hypothesis of this experiment was that trait mindfulness would moderate the stress response. In support of this, we found that those with higher trait mindfulness reported smaller increases in self-reported stress and cortisol concentration (Lin et al., in revision). This finding is in line with numerous reports in the literature suggesting that mindfulness has a buffering effect on the effects of both acute (Brown et al., 2012; Manigault et al., 2018) and chronic stress (Britton et al., 2012). Furthermore, mindfulness-based treatments have a clear effect on reducing self-reported stress in healthy individuals (Khoury et al., 2015). While the psychological and physiological mechanisms responsible for this stress reduction have not been fully elucidated, monitoring and acceptance (Ede et al., 2020), affect labelling (Lin et al., in revision), reduced amygdala connectivity (Bauer et al., 2019) and heart-rate variability (Watford et al., 2020) have been suggested as potential candidates in this pathway.

In previous work (Lim et al., 2018), we reported that trait mindfulness was associated with a third named dynamic connectivity state: the task-ready state. In that study, we divided participants into those high and low in trait mindfulness, and found that individuals high on trait mindfulness spend relatively more time in the TRS in a subsequent resting-state scan (Wong et al., 2018). With this finding in mind, we tested whether the significant decrease in TRS due to the TSST had any relationship to self-reported mindfulness measured using the Five Facet Mindfulness Questionnaire (FFMQ) in the current data set. Mindfulness scores were not correlated with the proportion of time spent in the TRS at baseline. Instead, we found that higher mindfulness scores were correlated with smaller reductions in TRS following stress.

An important point to appreciate is that the relationship between mindfulness and the TRS in this experiment was only uncovered after perturbation by stress. This belies our previous claim that the TRS encodes a constant, trait-like propensity to be mindful, and instead suggests that it is a functional configuration that is less vulnerable in mindful individuals during systemic challenge. In our previous report, we posited that the TRS may play a role in supporting cognitive flexibility (Lim et al., 2018), and this module is known to be impaired by stress, particularly in men (Shields et al., 2016). While we did not measure cognitive flexibility in this paradigm, our current results provide additional impetus to formally test this hypothesis.

Interestingly, changes in TRS correlated with an objective marker of stress (cortisol AUCg) but not with self-report, even though changes in self-reported stress and cortisol output themselves were correlated. Using the TSST, Reinelt et al. (2019) reported correlations between change in eigenvector centrality in the thalamus and self-reported stress (and non-significant associations of thalamic centrality with cortisol), suggesting that information about subjective state may be carried in the ascending relay of sensory signals rather than at the cortical level. Further, our data suggest that dynamic functional connectivity in the cortex may contain information about the objective effects of stress that is not consciously accessible to individuals.

The majority of studies of mindfulness using resting-state activity have focused on finding the static (Bilevicius et al., 2018) or dynamic (Mooneyham et al., 2017; Lim et al., 2018) connectivity markers of trait mindfulness, or changes in functional connectivity following mindfulness training (Brewer et al., 2011). While the results from these studies are somewhat mixed, some consensus has emerged that the insula is a key node associated with mindfulness (Falcone and Jerram, 2018), and that the default mode network is also likely to play a critical role. The distinguishing features of the TRS show good concordance with this extant literature, with strong within-network correlations in the default-mode network and salience network, and anti-correlations between the default mode network and key nodes of this network, including precentral gyrus, the anterior cingulate cortex and the insula.

### Functional connectivity does not return to baseline within 30 minutes of stress

We collected a third resting-state fMRI scan at 30 minutes after the TSST to test whether dynamic functional connectivity would return to baseline after recovery from stress. We found no significant differences between RS_2_ and RS_3_: TRS remained significantly lower in RS_3_, and LAS continued its upward trajectory, with significantly greater levels than in RS_1_. These data are in line with studies and showing a relatively slow return of connectivity to baseline levels following acute stress (Reinelt et al., 2019). In agreement with a model proposed by Hermans et al. (2014), this time course of change is more similar to the trajectory of cortisol release, which only recovered to baseline at the conclusion of the experiment (T6), than self-reported stress, which was already significantly lower before RS_3_ (and comparable to baseline levels) than before RS_2_. Recalling that TRS changes were associated with cortisol AUCg, this lack of recovery to baseline in RS_3_ is a further indication that functional connectivity changes seen as a result of stress are more strongly related to objective markers than with self-report.

### The arousal states are consistently coupled

In both this experiment and in previous data (Teng et al., 2019), we found that changes in HAS and LAS across state (stress and sleep deprivation) were highly negatively correlated, suggesting a tight coupling between these states even with changes in physiology and behaviour. Interestingly, this oppositional relationship was observed even as the proportion of time spent in both the HAS and LAS increased numerically after the TSST. These data solidify our belief that the two states operate in tandem to govern brain function (Wang et al., 2016), although as noted above, it is less clear from this dataset that arousal levels are the exclusive output of this balance.

## Conclusion

In summary, we have built upon previous work exploring the functional significance of three brain connectivity states to construct a model linking trait mindfulness, stress reactivity, and brain function. These findings further our understanding of how mindfulness moderates the stress response, and also hone our definition of how dynamic connectivity is altered by arousal and stress.

## Materials and Methods

### Participants

41 male participants were recruited from the National University of Singapore through online advertisements and word-of-mouth. We excluded females from this study to eliminate sex differences in response to stress, and the potential influence of the menstrual cycle and oral contraceptive use. One participant was excluded from the study for non-compliance to the protocol instructions, resulting in a final sample size of 40 (mean age (sd) = 23.03 (3.23)). All participants were screened to ensure that they had no history of long-term physical or psychological disorders, for right-handedness (Oldfield, 1971), and for normal or corrected-to-normal vision. Participants were only admitted to the study if they had no prior knowledge of the study protocol, or exposure to similar social stress studies. Finally, they were excluded if they reported any contraindications for MRI scanning. The study was approved by the National University of Singapore Institutional Review Board. All participants provided written informed consent, and were reimbursed with SGD $50 for their participation.

### Study Protocol

All testing sessions were conducted between 14:00 pm and 17:00 pm to account for diurnal fluctuations of cortisol (Nogeire et al., 1971). Participants first completed a 20-minute computerized cognitive task (results not reported in this communication), followed by a 10-minute eyes-open resting-state fMRI (RS_1_). Participants then performed the Trier Social Stress Test (TSST; see below) as a stress manipulation. Thereafter, participants underwent 2 additional resting-state fMRI scans (RS_2_ and RS_3_), interspaced with a 5-minute high-resolution MPRAGE structural scan. After the last MRI scan, participants remained in the lab for a 30-minute recovery period, before they were debriefed and reimbursed for their time.

Throughout the study, salivary cortisol and subjective stress ratings were collected at specific times during the protocol. The time points for all major data acquisition milestones were dictated by a pre-determined time schedule that research assistants adhered to as closely as possible; Figure 1 shows this experimental protocol with timing relative to the start of the TSST, together with the actual acquisition timelines of each individual participant.

### Stress Manipulation

The Trier Social Stress Test (TSST) is a common laboratory-administered protocol to induce psychosocial stress (Kirschbaum et al., 1993). At the start of the test, a research assistant informed the participant that they had to prepare a speech for a mock job interview in front of two “behavioural experts”. They were told that their performance would be video recorded and analysed. Participants were then left alone in a room for 5 minutes with writing material. They were allowed to make notes during this time, but these were collected from them at the end of the preparation period. Participants were then ushered by the research assistant to an adjacent room and introduced to the evaluation panel, which was always comprised of one male and one female confederate. Panellists were trained to withhold any social feedback and to maintain a neutral disposition at all times. We positioned a camera in plain sight of the participant that appeared to be operational, although no actual video recording was conducted. The participant was cued to begin their prepared speech, which was terminated after 5 minutes. If they fell silent or ran out of material before the end of the 5-minute period, one of the panellists prompted the participant to continue by selecting an appropriate question from a predetermined list. In the second 5-minute interview period, participants were then made to perform an unexpected mental arithmetic task. They were told to subtract 17 repeatedly from 2023 until they reach zero; to further induce stress, participants were urged to pick up their pace during long pauses, and had to restart their count if mistakes were made.

### Salivary Cortisol

To assess stress reactivity, we collected salivary cortisol using Salivette (Sarstedt, Nümbrecht, Germany) at five time points (Figure 1). During each cortisol collection, participants were instructed to place the cotton swab in their mouth for 2 minutes and told to avoid contamination with their hands. All salivettes were stored at −75°C before analysis in a commercial biotechnology lab. Saliva samples were centrifuged at 20°C (1000 x g, 2 minutes) and analyzed using salivary cortisol immunoassay (IBL International GMBH, Hamburg, Germany) with a sensitivity of 0.005 µg/dL. The intra and inter-assay coefficients of variation were all under 4%. The cortisol values were positively skewed and were thus log transformed for statistical analyses. We calculated two measures of cortisol output using the trapezoidal integration method of Pruessner et al. (2003). Area under the cortisol curve with respect to ground (AUCg) was defined as the total cortisol concentration over the duration of the experiment. Cortisol concentration increase related specifically to the stress induction (AUCi) was calculated by subtracting baseline production from AUCg. One subject did not produce adequate saliva sample for the immunoassay, and was not included in cortisol analysis.

### Subjective mood ratings

Subjectively experienced mood was measured at six time points (Figure 1), typically at the same time as cortisol collection. Participants indicated their immediate levels of Stress, Energy and Alertness on a nine point Likert scale from 0 = *not all* to 9 = *extremely*. For the remainder of the manuscript, we focus analysis on Stress and Alertness at only three of these points (T2-4).

### Self-reported mindfulness

We measured dispositional mindfulness using the Five Facet Mindfulness Questionnaire (FFMQ; Baer et al., 2006). The 39-item inventory evaluates five distinct components of mindfulness: Observing, Describing, Non-Reactivity, Non-Judging, and Acting with Awareness. Items were rated from 1 = *never or very rarely true* to 5 = *very often or always true*, and reversed scored for Non-Judging and Acting with Awareness subscales. The questionnaire showed good internal consistency with a Cronbach’s alpha coefficient of 0.85, and as such, we used full-scale FFMQ scores for analysis.

### fMRI Acquisition

Resting-state fMRI scans were collected on a 3-Tesla Siemens PrismaFit system (Siemens, Erlangen, Germany) using an interleaved gradient echo-planar imaging sequence (TR = 2000 ms, TE = 30 ms, flip angle = 90°, field-of-view = 192 × 192 mm, voxel size = 3 × 3 × 3 mm). 36 oblique axial slices were obtained, and 300 volumes (10 minutes) were collected for each scan. High-resolution structural images were collected using an MPRAGE sequence (TR = 2300 ms, TI = 900 ms, FA = 8°, voxel size = 1 × 1 × 1 mm, FOV = 256 × 240 mm, 192 slices).

Participants were instructed to remain still and keep their eyes open during the resting-state scan while not thinking about anything in particular. To ensure compliance, concurrent eye videos were acquired using an MR compatible camera (12M-I eye-tracking camera; MRC Systems GmbH, Germany) placed over the right eye. Pre-recorded reminders (e.g., “Open your eyes.”) were delivered whenever participants closed their eyes for more than 10 s. This procedure was previously used (Yeo et al. 2015) to ensure that participants kept their eyes open during resting-state scans, as this can affect the results of connectivity analysis (Yan et al., 2009).

### Resting-state fMRI analysis

Resting-state scans were preprocessed in accordance to the previously described procedure in Yeo et al. (2015), using a combination of FSL (Smith et al., 2004; Jenkinson et al., 2012), SPM (Wellcome Department of Cognitive Neurology, London, UK), and FreeSurfer (http://surfer.nmr.mgh.harvard.edu; Fischl, 2012). Briefly, preprocessing steps involved (i) discarding the first four frames of each run, (ii) slice time correction, (iii) head-motion correction using rigid body translation and rotation parameters, (iv) functional and structural images were aligned using Boundary-Based Registration following FreeSurfer surface reconstruction. Whole brain, white matter and ventricular masks were then defined based on structural segmentation, then transformed to subject space. White matter segmentation was performed with 1-voxel erosion. (v) Linear trend removal was subsequently performed, with bandpass temporal filtering (0.009 – 0.08 Hz), and linear regression of spurious signal (head motion, whole brain signal, white matter signal, ventricle signal, and their derivatives). (vi) Functional data of individual subjects were then projected onto MNI-152 space, downsampled to 2 mm voxels and then smoothed with a 6-mm full width half maximum kernel.

Global signal regression was carried out as a part of the preprocessing pipeline to remove potential nuisance components in the data (Liu et al., 2017; McAvoy et al., 2019). Global signal power, or the standard deviation of the average percentage change in the signal time course of the whole brain (Wong et al., 2013), was subsequently calculated

Head motion was calculated based on two measures: framewise displacement (FD) and variance of temporal derivative of time courses over voxels (DVARS; Power et al. 2012). Volumes having FD > 0.2 mm or DVARS >5% were marked as high motion. As we intended to perform dynamic functional connectivity analysis, motion scrubbing – or the removal of high motion volumes – was not conducted as this removal can have an impact on the temporal pattern of the underlying functional connectivity (Power et al., 2012). Instead, one volume before and two volumes after each high motion volume were also marked, and these frames were interpolated from surrounding data. No subject was excluded from the analysis for having more than 50% of total volumes marked as high motion.

### Dynamic functional connectivity analysis

Dynamic functional connectivity analysis was performed using the multiplication of temporal derivatives method described by Shine et al. (2015). 114 cortical ROIs were first extracted from the 17-network parcellation by Yeo et al. (2011). The coupling between each pairwise set of 114 ROIs was then estimated by multiplying the first-derivatives of the averaged BOLD time series. Connectivity at each time point was then estimated by computing a simple moving average of the multiplied temporal derivative time course using the recommended window size of 7 TRs, for a total of 292 coupling matrices per participant, each containing 6441 (114 × 113/2) unique coupling values.

Coupling matrices were than concatenated across the 40 participants for all three resting-state fMRI scans, and k-means clustering was performed to classify each matrix using squared Euclidean distance as the cost function. We used a k = 5 solution, for consistency with our previous reports, and as recent work using a large (N = 7,500) dataset of resting-state scans suggests that this is an optimal number of clusters (Abrol et al., 2017). To confirm that our centroids were consistent with those obtained from previous analyses reported by our group (Lim et al., 2018; Patanaik et al., 2018), we performed Spearman’s correlations between the two sets of centroids. We then calculated the proportion of the run spent in each dynamic connectivity state.

### Statistical Analysis

All statistical analyses were conducted on SPSS 25.0 for Windows (IBM Corp., Armonk, N.Y., USA), and statistical significance for all analysis was set at α = 0.05. Change scores of our primary outcome variables are calculated by subtracting scores pre-TSST from post-TSST scores. Changes in proportion of time spent in dynamic connectivity state across the three resting-state fMRI scans were similarly calculated (RS_2_-RS_1_, RS_3_-RS_2_).

Paired samples t-tests were computed for proportion of time spent in each connectivity stage across all three resting-state fMRI scans, as well as self-reported stress, and salivary cortisol before and after the TSST.

Pearson’s correlation was performed between the change in self-reported arousal and stress, and the change in the proportion of time spent in the HAS and LAS from before to after the TSST. Spearman’s correlation was performed to determine the similarity between dynamic connectivity states found in this study with previously found states.

## Acknowledgements

We acknowledge the assistance of Jia Lin, Zaven Leow, TeYang Lau, Ksenia Vinogradova and Kamalakanna Mayalagu Vijayakumar in data collection. This study was supported by the National Medical Research Council, Singapore (STaR/0015/2013), the National Research Foundation Science of Learning grant (NRF2016-SOL002-001), and by start-up funding from Duke-NUS Medical School. The funding agencies had no role in the conduct of the research or preparation of the article.

## Data availability

Data reported in this study are available at https://osf.io/6kr34/

## References

Abrol A, Damaraju E, Miller RL, Stephen JM, Claus ED, Mayer AR, Calhoun VD (2017) Replicability of time-varying connectivity patterns in large resting state fMRI samples. Neuroimage 163:160–176.

Allen EA, Damaraju E, Plis SM, Erhardt EB, Eichele T, Calhoun VD (2014) Tracking whole-brain connectivity dynamics in the resting state. Cerebral Cortex 24.3:663–76.

Baer RA, Smith GT, Hopkins J, Krietemeyer J, Toney L (2006) Using Self-Report Assessment Methods to Explore Facets of Mindfulness. Assessment 13:27–45.

Barttfeld P, Petroni A, Baez S, Urquina H, Sigman M, Cetkovich M, Torralva T, Torrente F, Lischinsky A, Castellanos X, Manes F, Ibanez A (2014) Functional connectivity and temporal variability of brain connections in adults with attention deficit/hyperactivity disorder and bipolar disorder. Neuropsychobiology 69:65–75.

Bauer CCC, Caballero C, Scherer E, West MR, Mrazek MD, Phillips DT, Whitfield-Gabrieli S, Gabrieli JDE (2019) Mindfulness training reduces stress and amygdala reactivity to fearful faces in middle-school children. Behav Neurosci 133:569–585.

Bilevicius E, Smith SD, Kornelsen J (2018) Resting-State Network Functional Connectivity Patterns Associated with the Mindful Attention Awareness Scale. Brain Connect 8:40–48.

Brewer JA, Mallik S, Babuscio TA, Nich C, Johnson HE, Deleone CM, Minnix-Cotton CA, Byrne SA, Kober H, Weinstein AJ, Carroll KM, Rounsaville BJ (2011) Mindfulness training for smoking cessation: results from a randomized controlled trial. Drug Alcohol Depend 119:72–80.

Britton WB, Shahar B, Szepsenwol O, Jacobs WJ (2012) Mindfulness-based cognitive therapy improves emotional reactivity to social stress: results from a randomized controlled trial. Behav Ther 43:365–380.

Brown KW, Ryan RM (2003) The benefits of being present: mindfulness and its role in psychological well-being. J Pers Soc Psychol 84:822–848.

Brown KW, Weinstein N, Creswell JD (2012) Trait mindfulness modulates neuroendocrine and affective responses to social evaluative threat. Psychoneuroendocrinology 37:2037–2041.

Chang C, Leopold DA, Scholvinck ML, Mandelkow H, Picchioni D, Liu X, Ye FQ, Turchi JN, Duyn JH (2016) Tracking brain arousal fluctuations with fMRI. Proc Natl Acad Sci USA 113:4518–4523.

Chang J, Yu R (2019) Hippocampal connectivity in the aftermath of acute social stress. Neurobiology of Stress 11:100195.

Dickerson SS, Kemeny, ME (2004) Acute stressors and cortisol responses: a theoretical integration and synthesis of laboratory research. Psychological bulletin, 130.3:355.

Dinges DF (1995) An overview of sleepiness and accidents. J Sleep Res 4:4–14.

Doll A, Holzel BK, Boucard CC, Wohlschlager AM, Sorg C (2015) Mindfulness is associated with intrinsic functional connectivity between default mode and salience networks. Front Hum Neurosci 9:461.

Ede DE, Walter FA, Hughes JW (2020) Exploring How Trait Mindfulness Relates to Perceived Stress and Cardiovascular Reactivity. Int J Behav Med.

Falahpour M, Chang C, Wong CW, Liu TT (2018) Template-based prediction of vigilance fluctuations in resting-state fMRI. Neuroimage 174:317–327.

Falcone G, Jerram M (2018) Brain activity in mindfulness depends on experience: a meta-analysis of fMRI studies. Mindfulness 9:1319–1329.

Fischl B (2012) FreeSurfer. Neuroimage 62:774–781.

Gotink RA, Meijboom R, Vernooij MW, Smits M, Hunink MG (2016) 8-week Mindfulness Based Stress Reduction induces brain changes similar to traditional long-term meditation practice - A systematic review. Brain Cogn 108:32–41.

Hermans EJ, Henckens MJ, Joels M, Fernandez G (2014) Dynamic adaptation of large-scale brain networks in response to acute stressors. Trends Neurosci 37:304–314.

Jenkinson M, Beckmann CF, Behrens TE, Woolrich MW, Smith SM (2012) Fsl. Neuroimage 62:782–790.

Kabat-Zinn J (1991) Full catastrophe living: Using the wisdom of your body and mind to face stress, pain, and illness. New York, N.Y.: Dell Pub., a division of Bantam Doubleday Dell Pub. Group.

Khoury B, Sharma M, Rush SE, Fournier C (2015) Mindfulness-based stress reduction for healthy individuals: A meta-analysis. J Psychosom Res 78:519–528.

Kilpatrick LA, Suyenobu BY, Smith SR, Bueller JA, Goodman T, Creswell JD, Tillisch K, Mayer EA, Naliboff BD (2011) Impact of Mindfulness-Based Stress Reduction training on intrinsic brain connectivity. Neuroimage 56:290–298.

Kirschbaum C, Pirke KM, Hellhammer DH (1993) The ‘Trier Social Stress Test’--a tool for investigating psychobiological stress responses in a laboratory setting. Neuropsychobiology 28:76–81.

Kudielka BM, Hellhammer DH, Kirschbaum C, (2007) Ten years of research with the trier social stress test revisited. In: Harmon-Jones E, Winkielman P, editor. Social Neuroscience: Integrating Biological and Psychological Explanations of Social Behavior. Guilford Press, New York. pp. 56–83.

Lim J, Teng J, Patanaik A, Tandi J, Massar SAA (2018) Dynamic functional connectivity markers of objective trait mindfulness. Neuroimage 176:193–202.

Lin J, Massar SAA, Lim J (in revision) Trait mindfulness moderates reactivity to social stress in an all-male sample. Mindfulness.

Liu TT, Nalci A, Falahpour M (2017) The global signal in fMRI: Nuisance or Information? Neuroimage 150:213–229.

Manigault AW, Woody A, Zoccola PM, Dickerson SS (2018) Trait mindfulness predicts the presence but not the magnitude of cortisol responses to acute stress. Psychoneuroendocrinology 90:29–34.

Marusak HA, Elrahal F, Peters CA, Kundu P, Lombardo MV, Calhoun VD, Goldberg EK, Cohen C, Taub JW, Rabinak CA (2018) Mindfulness and dynamic functional neural connectivity in children and adolescents. Behav Brain Res 336:211–218.

Massar SAA, Liu JCJ, Mohammad NB, Chee MWL (2017) Poor habitual sleep efficiency is associated with increased cardiovascular and cortisol stress reactivity in men. Psychoneuroendocrinology 81:151–156.

McAvoy MP, Tagliazucchi E, Laufs H, Raichle ME (2019) Human non-REM sleep and the mean global BOLD signal. J Cereb Blood Flow Metab 39:2210–2222.

McEwen BS (2006) Sleep deprivation as a neurobiologic and physiologic stressor: Allostasis and allostatic load. Metabolism. 55.10:S20–3.

Minkel J, Moreta M, Muto J, Htaik O, Jones C, Basner M, Dinges D (2014) Sleep deprivation potentiates HPA axis stress reactivity in healthy adults. Health Psychol 33(11):1430–4.

Mooneyham BW, Mrazek MD, Mrazek AJ, Schooler JW (2016) Signal or noise: brain network interactions underlying the experience and training of mindfulness. Ann N Y Acad Sci 1369:240–256.

Mooneyham BW, Mrazek MD, Mrazek AJ, Mrazek KL, Phillips DT, Schooler JW (2017) States of Mind: Characterizing the Neural Bases of Focus and Mind-wandering through Dynamic Functional Connectivity. J Cogn Neurosci 29:495–506.

Nogeire C, Fukushima D, Weitzman ED, Roffwarg H, Hellman L, Gallagher TF (1971) Twenty-four Hour Pattern of the Episodic Secretion of Cortisol in Normal Subjects. The Journal of Clinical Endocrinology & Metabolism 33:14–22.

Oldfield RC (1971) The assessment and analysis of handedness: the Edinburgh inventory. Neuropsychologia 9:97–113.

Parkinson TD, Kornelsen J, Smith SD (2019) Trait Mindfulness and Functional Connectivity in Cognitive and Attentional Resting State Networks. Front Hum Neurosci 13:112.

Patanaik A, Tandi J, Ong JL, Wang C, Zhou J, Chee MWL (2018) Dynamic functional connectivity and its behavioral correlates beyond vigilance. Neuroimage 177:1–10.

Posner MI, Petersen SE (1990) The attention system of the human brain. Annu Rev Neurosci 13:25–42.

Power JD, Barnes KA, Snyder AZ, Schlaggar BL, Petersen SE (2012) Spurious but systematic correlations in functional connectivity MRI networks arise from subject motion. Neuroimage 59:2142–2154.

Pruessner JC, Kirschbaum C, Meinlschmid G, Hellhammer DH (2003) Two formulas for computation of the area under the curve represent measures of total hormone concentration versus time-dependent change. Psychoneuroendocrinology 28:916–931.

Reinelt J, Uhlig M, Muller K, Lauckner ME, Kumral D, Schaare HL, Baczkowski BM, Babayan A, Erbey M, Roebbig J, Reiter A, Bae YJ, Kratzsch J, Thiery J, Hendler T, Villringer A, Gaebler M (2019) Acute psychosocial stress alters thalamic network centrality. NeuroImage 199:680–690.

Sarter M, Givens B, Bruno JP (2001) The cognitive neuroscience of sustained attention: where top-down meets bottom-up. Brain Res Brain Res Rev 35:146–160.

Shields GS, Trainor BC, Lam JC, Yonelinas AP (2016) Acute stress impairs cognitive flexibility in men, not women. Stress 19:542–546.

Shine JM, Koyejo O, Bell PT, Gorgolewski KJ, Gilat M, Poldrack RA (2015) Estimation of dynamic functional connectivity using Multiplication of Temporal Derivatives. Neuroimage 122:399–407.

Smith SM, Jenkinson M, Woolrich MW, Beckmann CF, Behrens TE, Johansen-Berg H, Bannister PR, De Luca M, Drobnjak I, Flitney DE, Niazy RK, Saunders J, Vickers J, Zhang Y, De Stefano N, Brady JM, Matthews PM (2004) Advances in functional and structural MR image analysis and implementation as FSL. Neuroimage 23 Suppl 1:S208–219.

Spiegel K, Leproult R, Van Cauter E (1999) Impact of sleep debt on metabolic and endocrine function. Lancet 354(9188):1435–9.

Teng J, Ong JL, Patanaik A, Tandi J, Zhou JH, Chee MWL, Lim J (2019) Vigilance declines following sleep deprivation are associated with two previously identified dynamic connectivity states. Neuroimage 200:382–390.

Turchi J, Chang C, Ye FQ, Russ BE, Yu DK, Cortes CR, Monosov IE, Duyn JH, Leopold DA (2018) The Basal Forebrain Regulates Global Resting-State fMRI Fluctuations. Neuron 97:940–952 e944.

Veer IM, Oei NY, Spinhoven P, van Buchem MA, Elzinga BM, Rombouts SA (2011) Beyond acute social stress: increased functional connectivity between amygdala and cortical midline structures. NeuroImage 57.4:1534–41.

Wang C, Ong JL, Patanaik A, Zhou J, Chee MW (2016) Spontaneous eyelid closures link vigilance fluctuation with fMRI dynamic connectivity states. Proc Natl Acad Sci U S A 113:9653–9658.

Watford TS, O’Brien WH, Koerten HR, Bogusch LM, Moeller MT, Sonia Singh R, Sims TE (2020) The mindful attention and awareness scale is associated with lower levels of high-frequency heart rate variability in a laboratory context. Psychophysiology 57:e13506.

Wong CW, Olafsson V, Tal O, Liu TT (2013) The amplitude of the resting-state fMRI global signal is related to EEG vigilance measures. Neuroimage 83:983–990.

Wong KF, Massar SAA, Chee MWL, Lim J (2018) Towards an objective measure of mindfulness: replicating and extending the features of the breath-counting task. Mindfulness.

Yeo BT, Tandi J, Chee MW (2015) Functional connectivity during rested wakefulness predicts vulnerability to sleep deprivation. Neuroimage 111:147–158.

Yeo BT, Krienen FM, Sepulcre J, Sabuncu MR, Lashkari D, Hollinshead M, Roffman JL, Smoller JW, Zollei L, Polimeni JR, Fischl B, Liu H, Buckner RL (2011) The organization of the human cerebral cortex estimated by intrinsic functional connectivity. Journal of neurophysiology 106:1125–1165.

Zhang X, Huettel SA, O’Dhaniel A, Guo H, Wang L (2019) Exploring common changes after acute mental stress and acute tryptophan depletion: Resting-state fMRI studies. Journal of psychiatric research 113:172–80.

