## Supplemental Figure 1 for "Inter-relationships between changes in stress, mindfulness, and dynamic functional connectivity in response to a social stressor"

Supplemental information

Reproducibility of dynamic connectivity states

In several previous reports, we have referred to three named dynamic connectivity states, the “high arousal state”, the “low arousal state”, and the “task-ready state” (Wang et al., 2018, Lim et al., 2018, Patanaik et al., 2019, Teng et al., 2019). We have claimed that the centroids for these states are reproducible when unsupervised (k-means) clustering is used in independent datasets. While the states are easily distinguishable on visual inspection, and show high correlations across datasets, inspection of the full correlation matrix between datasets reveals that there is not an optimized solution where each state has one and only one best-matched counterpart; specifically, the state previously identified as the TRS in Lim et al. (2018) is more highly correlated with the unnamed "State 5" in the current experiment than the state that is identifiable as "TRS" (see Supplementary Figure 1a).

To address this ambiguity, we attempted to create more stable estimates of the DCS centroids by increasing the amount of data used. Four separate datasets were compiled, consisting of 122 undergraduate participants (50 males; mean (sd) age = 22.8 (2.91)). Subjects from two of the datasets provided two independent resting state scans, resulting in a total of 173 scans entering the canonical analysis. The four separate datasets were preprocessed and analysed in an identical manner as described in Materials and Methods. Unsupervised k-means clustering was performed on the resulting datasets to obtain the 5 "canonical" centroids. Using these, we clearly show that across both the current dataset as well as that of Lim et al. (2018), all ambiguous state pairings are resolved, and an optimal one-to-one match of states can be found (as can be seen on the diagonal of the matrix of Supplementary Figure 1b and c). This analysis suggests that the states under study are unique and reproducible across different samples.

We have made the connectivity matrices of these canonical centroids freely available on our github at (https://github.com/awakelab/Dynamic-Functional-Connectivity-MTD-)

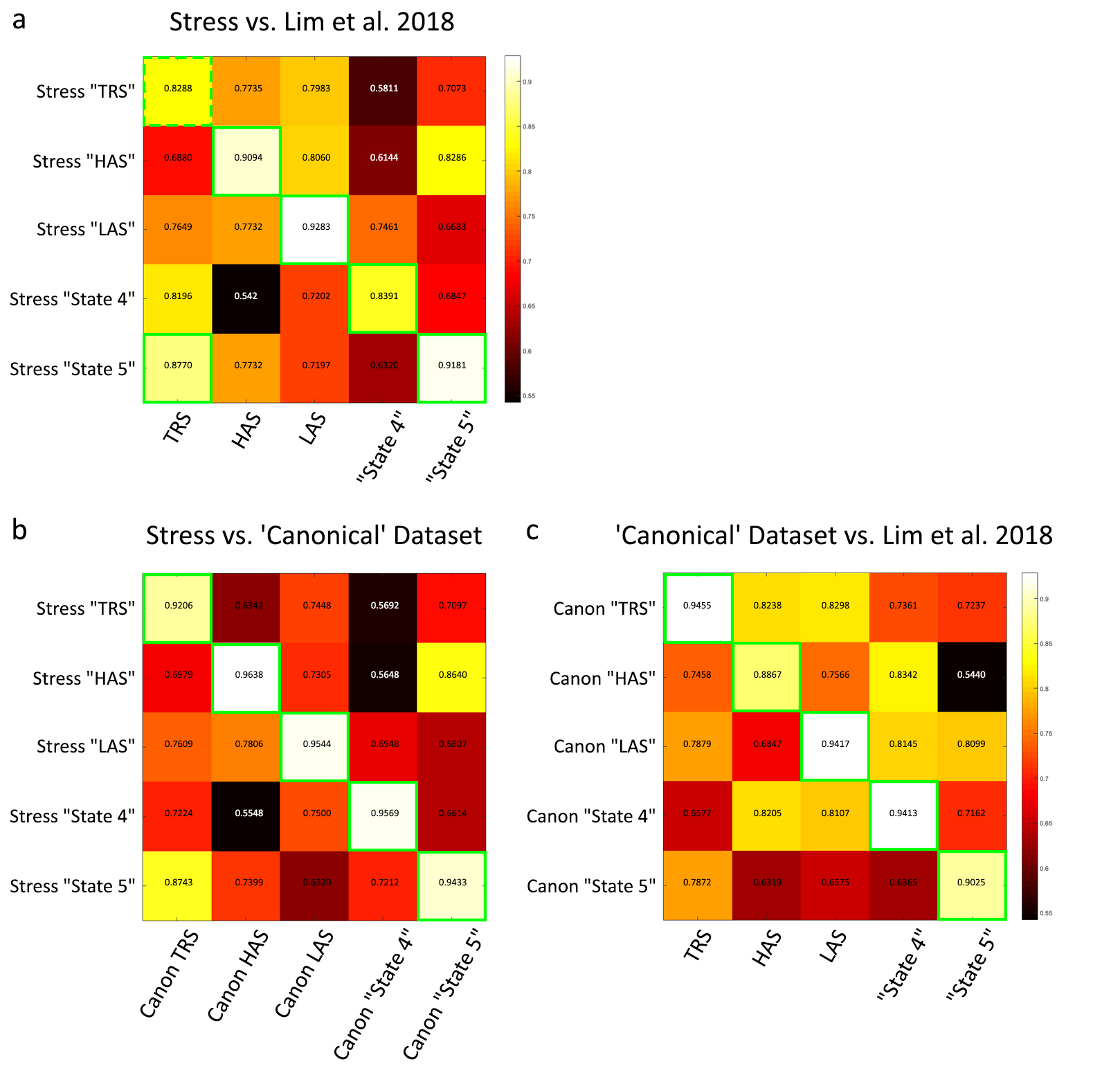

**Supplementary Figure 1**. Comparisons between centroids obtained in three datasets using spearman's correlation. Centroids from the current dataset is compared with a) a previously published dataset (Lim et al., 2018), and b) a compilation dataset used to create "canonical" centroids. A further comparison was shown c) between the Lim et al. (2018) dataset with the "canonical" centroids.
